## Supplementary information for "Transcriptomic signatures of brain regional vulnerability to Parkinson’s disease"

### Supplementary Tables

**Supplementary Table 1 Number of samples for each Braak stage-related region across the six donors in AHBA.** Each one of the Braak regions is represented by one or multiple anatomical structures; names are according to the ontology of AHBA.

| Braak stage-related regions | Anatomical structures | Donor 9861 | Donor 10021 | Donor 12876 | Donor 14380 | Donor 15496 | Donor 15697 | Total |
|---|---|---|---|---|---|---|---|---|
| R1 | Myelencephalon | 76 | 85 | 16 | 44 | 35 | 23 | 279 |
| R2 | Pontine tegmentum | 40 | 38 | 2 | 19 | 21 | 21 | 141 |
| R3 | Substantia nigra, basal nucleus of meynert, right, basal nucleus of meynert, left, CA2 field | 20 | 25 | 7 | 12 | 13 | 12 | 89 |
| R4 | Amygdala, occipito-temporal gyrus | 25 | 29 | 13 | 13 | 13 | 14 | 107 |
| R5 | Cingulate gyrus, temporal lobe | 178 | 117 | 62 | 97 | 76 | 88 | 618 |
| R6 | Frontal lobe, parietal lobe | 242 | 192 | 84 | 112 | 96 | 101 | 827 |

**Supplementary Table 2**
List of all BRGs with Braak correlations, fold-change between Braak region 1 and 6, and its Benjamini-Hochberg-corrected p-values: *Supplementary Table 2.xlsx*.

**Supplementary Table 3**
Functional GO-terms associated with negative correlated BRGs: *Supplementary Table 3.xlsx*.

**Supplementary Table 4**
Functional GO-terms associated with positive correlated BRGs: *Supplementary Table 4.xlsx*.

**Supplementary Table 5 Number of samples in PD microarray dataset.** Braak stage-related regions are represented by one anatomical structure.

| Braak stage-related regions | Anatomical structures | PD | iLBD | Controls |
|---|---|---|---|---|
| R1 | Medulla oblongata (Myelencephalon) | 10 | 11 | 5 |
| R2 | Locus ceruleus | 17 | 12 | 9 |
| R3 | Substantia nigra | 12 | 8 | 8 |

**Supplementary Table 6**
Demographic and pathological data of postmortem brain tissue samples of controls, incidental Lewy body disease (iLBD), and Parkinson's disease (PD) cases in the PD microarray dataset: *Supplementary Table 6.xlsx*.

**Supplementary Table 7 Number of samples in PD RNA-seq dataset.** Braak stage-related regions are represented by one anatomical structure.

| Braak stage-related regions | Anatomical structures | PD | Controls |
|---|---|---|---|
| R3 | Substantia nigra | 10 | 11 |
| R4/R5 | Medial temporal gyrus | 5 | 9 |

**Supplementary Table 8**

Demographic and pathological data of postmortem brain tissue samples of controls and PD cases in the PD RNA-seq dataset: *Supplementary Table 8.xlsx*.

**Supplementary Table 9**

Modules with the number of genes, eigengene correlation with Braak stages, Benjamini-Hochberg-corrected P-values of correlations, and genes within the modules: *Supplementary Table 9.xlsx*.

**Supplementary Table 10 Literature overview on protective and toxic effects of α-synuclein.** Laboratory studies that examined the effect of wild-type or mutant α-synuclein using various experiments. A53T and A30P are familial Parkinson's disease pathogenic mutations. DA: dopamine, PD: Parkinson's disease.

| **Study** | **Samples** | **Molecular type of α-synuclein** | **Outcome** | **Wild-type α-synuclein is protective?** |
|---|---|---|---|---|
| Neystat et al. (1999)[1] | Substantia nigra and cortex from 9 PD patients and 8 controls | mRNA | Less α-synuclein mRNA was observed in the substantia nigra of PD patients implying that decreased levels may play a role in the pathogenesis. It is unlikely that this decrease simply reflects neuronal loss because a similar difference was observed after normalizing for *VMAT2* (a dopaminergic neuronal marker) expression. | Yes |
| Kholodilov et al. (1999)[2] | Rat model of induced programmed cell death in substantia nigra (striatal lesions with quinolinic acid) and mesencephalic DA neurons. | Protein, mRNA, DNA | α-Synuclein does not play in direct role in mediating cell death, but may play a role in compensatory or plasticity response in surviving neurons. | Yes |
| Abeliovich et al. (2000)[3] | Brains of wild-type and α-synuclein -/- mice. | Protein | Mutant mice display altered DA release, and reduced striatal DA and amphetamine-stimulated locomotion. Thus, α-synuclein is an essential presynaptic, activity-dependent negative regulator of DA neurotransmission. | Yes |
| Alves da Costa et al. (2000)[4] | Mouse neocortical neuronal cell line overexpressing wild-type or A53T mutant α-synuclein. | Protein | Wild-type α-synuclein exerts an antiapoptotic effect in neurons that appears to be abolished by the PD-related mutation. | Yes |
| Alves da Costa et al. (2002)[5] | Mouse neocortical neuronal cell line overexpressing wild-type α-synuclein. | Protein | α-Synuclein lowers the p53-dependent caspase 3 activation and natural toxin 6-hydroxydopamine abolishes the antiapoptotic phenotype by triggering α-synuclein aggregation. | Yes |
| Lee et al. (2001)[6] | Human teratocarcinoma and neuroblastoma cell lines overexpressing wild-type, A30P and A53T mutant α-synuclein. | Protein | Mutant α-synucleins increased levels of 8-hydroxyguanine, protein carbonyls, lipid peroxidation and 3-nitrotyrosine, and markedly accelerated cell death in response to all the insults examined. Wild-type α-synuclein is cell line- and insult-specific, suggesting both protective and non-protective properties. | Yes |
| Stefanis et al. (2001)[7] | Rat pheochromocytoma (dopaminergic) PC12 cell line overexpressing wild-type and A53T mutant α-synuclein. | Protein | Mutant α-synuclein enhances cell death, induces accumulation of autophagic–vesicular structures, and results in a loss of catecholamine storage granules and the capacity for depolarization-induced DA release. These effects may involve lysosomal and proteasomal degradation systems. | Yes |
| Hashimoto et al. (2002)[8] | Mouse hypothalamic tumor neuronal cell line overexpresssing α-synuclein, β-synuclein or control vector. | Protein, mRNA | Increased α-synuclein expression might protect cells from oxidative stress by inactivation of JNK via increased expression of JIP-1b/IB1. | Yes |
| Seo et al. (2002)[9] | Rat primary cortical and hippocampal neurons and microglia, and various neuronal cell lines. | Protein | At low concentrations, α-synuclein protects neurons against stress conditions through the PI3/Akt signaling pathway. At overexpression α-synuclein induced cytotoxicity. | Yes and no |

| | | | | |
|---|---|---|---|---|
| Cabin et al. (2002)[10] | Whole brain and hippocampal neurons from α-synuclein -/- and +/+ mice | Protein | α-Synuclein is required for the genesis, localization, and/or maintenance of at least some subset of vesicles that make up the reserve or resting pools of presynaptic vesicles | Yes |
| Wersinger et al. (2003)[11] | Ltk- fibroblasts, human neuroblastoma-derived SK-N-MC cells, and HEK 293 human embryonic kidney cells transfected with DAT and α-synuclein, and primary rat mesencephalic cultures. | Protein | α-Synuclein forms a complex with DAT in both rat brain tissues and transfected cells, which regulates the amount of DAT at the plasma membrane. In basal situations, α-synuclein tends to markedly decrease DA uptake by DAT as well as decrease DA-mediated oxidative stress and cell death. The primary function of presynaptic α-synuclein in dopaminergic neurons may be the regulation of DA homeostasis | Yes |
| Zourlidou et al. (2003)[12] | Neuronal ND7 cell line overexpressing wild-type, A30P and A53T mutant α-synuclein. | protein | Mutations in α-synuclein convert the protein from one that is able to modulate cell death into a protein with a damaging effect in neuronal cells. | Yes |
| Kingsbury et al. (2004)[13] | human substantia nigra and cortex of PD and control subjects. | mRNA | In substantia nigra neurons there seemed to be a negative correlation between cellular mRNA expression and disease duration. Lewy body formation is unlikely to be the result of overexpression of α-synuclein. | Yes |
| Chandra et al. (2004)[14] | α -/- β-/- and α -/- β+/+ synuclein mice. | Protein | Synucleins are not essential components of the basic machinery for neurotransmitter release but may contribute to the long-term regulation and or maintenance of presynaptic function. | Yes |
| Chandra et al. (2005)[15] | Mice that overexpress wildtype human α-synuclein, A30P mutant human α-synuclein, A53T mutant human α-synuclein, or wild-type mouse α-synuclein. | Protein | Transgenic expression of α-synuclein unexpectedly abolishes the lethal phenotype created by deletion of presynaptic chaperone protein CSPα in mice. | Yes |
| Oksman et al. (2006)[16] | α-Synuclein -/- and +/+ mice. | | A lack of α-synuclein sensitized the brain reward system, implying that the levels of α-synuclein expression may predispose an individual to drug abuse or to a number of psychiatric diseases. | Yes |
| Burré et al. (2010)[17] | Brains from wild-type mice, CSPα -/- mice, and CSPα -/- rescued by transgenic α-synuclein and HEK293 cells. | | Maintenance of continuous presynaptic SNARE-complex assembly required a non-classical chaperone activity mediated by synucleins. | Yes |
| Al-Wandi et al. (2010)[18] | 2 year old α-synuclein -/- mice and γ-synuclein -/- mice. | Protein | The absence of this protein makes striatal dopaminergic synapses more vulnerable to changes associated with aging of the nervous system. | Yes |
| Greten-Harrison et al. (2010) | Neuronal cultures of wildtype and αβγ-synuclein -/- mice. | Protein | Synuclein deficiency leads to altered synapse structure and physiology, age-dependent neuronal dysfunction, and impaired survival. | Yes |
| Anwar et al. (2011)[19] | αβγ-synuclein -/- mice. | Protein | Synucleins in some dopaminergic neurons limit synaptic neurotransmission through mechanisms that differ from those reported in other neurons, which have included vesicle pool redistribution or SNARE complex formation. | Yes |
| Beatman et al. (2016)[20] | α-synuclein -/-, -/+, +/+ mice, neuronal cells from strial and parietal cortical tissue. Brain tissue from patients with West Nile virus infection. | Protein | Native α-synuclein expression in neurons inhibits viral growth and injury in the CNS. | Yes |
| Chen et al. (2018)[21] | α-Synuclein +/- and -/- cell lines of dopaminergic neurons from human stem cells. | mRNA, protein | Reducing or completely removing *SNCA* alleles by CRISPR/Cas9n-mediated gene editing confers a measure of resistance to Lewy pathology. | No |

#### References


1. Neystat, M. *et al.* α-Synuclein Expression in Substantia Nigra and Cortex in Parkinson's Disease. *Mov. Disord.* **14**, 417–422 (1999).
2. Kholodilov, N. G. *et al.* Increased Expression of Rat Synuclein in the Substantia Nigra Pars

Compacta Identified by mRNA Differential Display in a Model of Developmental Target Injury. *J. Neurochem.* **73**, 2586–2599 (1999).
3. Abeliovich, A. *et al.* Mice Lacking α-Synuclein Display Functional Deficits in the Nigrostriatal Dopamine System. *Cell Neuron* **25**, 239–252 (2000).
4. Alves da Costa, C., Ancolio, K. & Checler, F. Wild-type but Not Parkinson's Disease-related Ala-53 Thr Mutant α-Synuclein Protects Neuronal Cells from Apoptotic Stimuli. *J. Biol. Chem.* **275**, 24065–24069 (2000).
5. Alves, C., Paitel, E. & Vincent, B. α-Synuclein Lowers p53-dependent Apoptotic Response of Neuronal Cells. *J. Biol. Chem.* **277**, 50980–50984 (2002).
6. Lee, M., Hyun, D., Halliwell, B. & Jenner, P. Effect of the overexpression of wild-type or mutant Neuronal Cells-synuclein on cell susceptibility to insult. *J. Neurochem.* **76**, 998–1009 (2001).
7. Stefanis, L., Larsen, K. E., Rideout, H. J., Sulzer, D. & Greene, L. A. Expression of A53T Mutant But Not Wild-Type Neuronal Cells-Synuclein in PC12 Cells Induces Alterations of the Ubiquitin-Dependent Degradation System, Loss of Dopamine Release, and Autophagic Cell Death. *J. Neurosci.* **21**, 9549–9560 (2001).
8. Hashimoto, M. *et al.* Neuronal Cells-Synuclein Protects against Oxidative Stress via Inactivation of the c-Jun N-terminal Kinase Stress-signaling Pathway in Neuronal Cells. *J. Biol. Chem.* **277**, 11465–11472 (2002).
9. Seo, J. *et al.* α-Synuclein regulates neuronal survival via Bcl-2 family expression and PI3/Akt kinase pathway. *FASEB J.* **16**, (2002).
10. Cabin, D. E. *et al.* Synaptic Vesicle Depletion Correlates with Attenuated Synaptic α-Synuclein. *J. Neurosci.* **22**, 8797–8807 (2002).
11. Wersinger, C., Prou, D., Vernier, P. & Sidhu, A. Modulation of dopamine transporter function by α-synuclein is altered by impairment of cell adhesion and by induction of oxidative stress. *FASEB J.* **17**, (2003).
12. Zourlidou, A., Smith, M. D. P. & Latchman, D. S. Modulation of cell death by α-synuclein is stimulus-dependent in mammalian cells. *Neurosci. Lett.* **340**, 234–238 (2003).
13. Kingsbury, A. E. *et al.* Alteration in α-Synuclein mRNA Expression in Parkinson's Disease. *Mov. Disord.* **19**, 162–170 (2004).
14. Chandra, S. *et al.* Double-knockout mice for α- and β-synucleins: Effect on synaptic functions. *Proc. Natl. Acad. Sci. U. S. A.* **101**, 14966–14971 (2004).
15. Chandra, S., Gallardo, G., Fernández-chacón, R., Schlüter, O. M. & Südhof, T. C. α-Synuclein Cooperates with CSPα in Preventing Neurodegeneration. *Cell* **123**, 383–396 (2005).
16. Oksman, M., Tanila, H. & Yavich, L. Brain reward in the absence of alpha-synuclein. *Mol. Neurosci.* **17**, 1191–1194 (2006).
17. Burré, J., Sharma, M., Tsetsenis, T., Buchman, V. & Südhof, T. C. α-Synuclein Promotes SNARE-Complex Assembly in vivo and in vitro. *Science* **329**, 1663–1667 (2010).
18. Al-wandi, A. *et al.* Absence of α-synuclein affects dopamine metabolism and synaptic markers in the striatum of aging mice. *Neurobiol. Aging* **31**, 796–804 (2010).
19. Anwar, S. *et al.* Functional Alterations to the Nigrostriatal System in Mice Lacking All Three Members of the Synuclein Family. *J. Neurosci.* **31**, 7264–7274 (2011).
20. Beatman, E. L. *et al.* Alpha-Synuclein Expression Restricts RNA Viral Infections in the Brain. *J. Virol.* **90**, 2767–2782 (2016).
21. Chen, Y. *et al.* Engineering synucleinopathy-resistant human dopaminergic neurons by CRISPR-mediated deletion of the *SNCA* gene. *Eur. J. Neurosci.* **49**, 510–524 (2019).

### Supplementary Figures

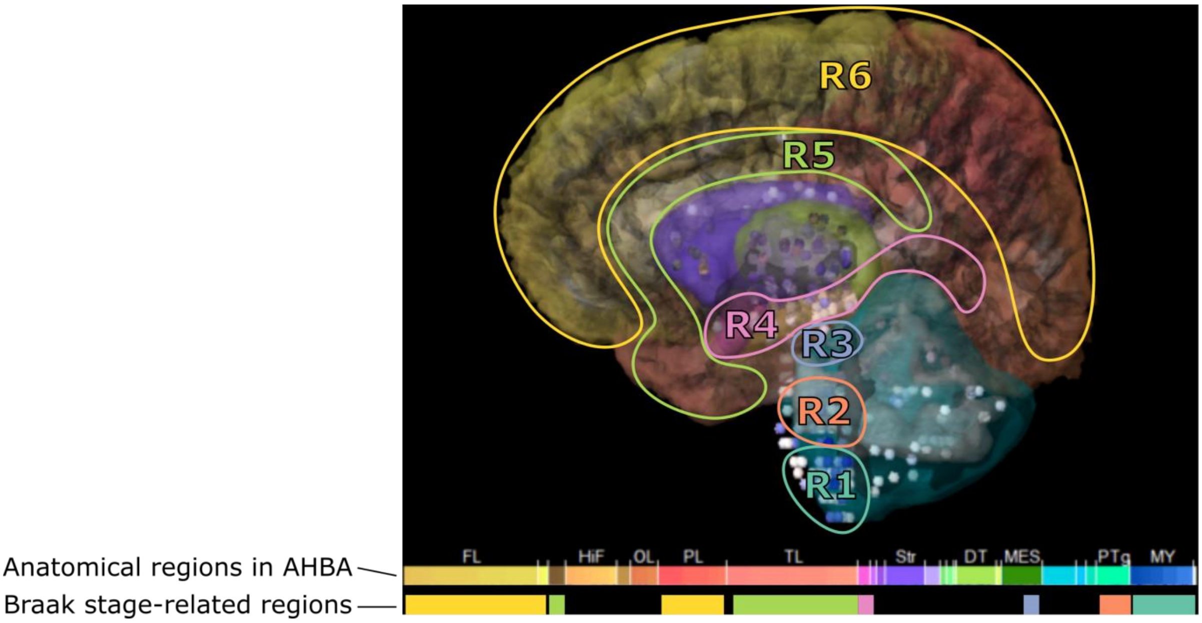


**Supplementary Figure 1 Defining Braak stage-related regions in the Allen Human Brain Atlas (AHBA).** Top row shows the anatomical regions and annotated colors as defined in the AHBA. Bottom row shows regions that are assigned as Braak stage-related regions where colors correspond to each one of the six regions R1-R6. See Supplementary Table 1 for names of anatomical structures and sample size information.

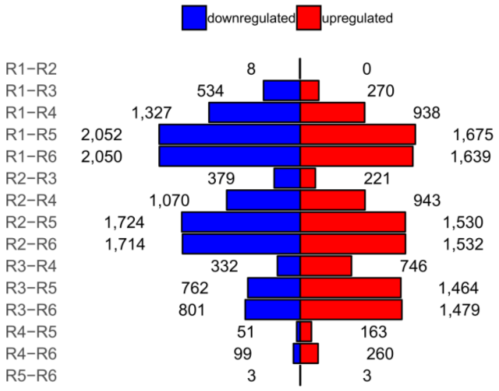


**Supplementary Figure 2 Number of differentially expressed genes between all pairs of six Braak stage-related regions R1-R6.** Differentially expressed genes were downregulated (blue) or upregulated (red) in the second region compared to the first region. Respectively, this supports a negative or positive correlation with Braak stages.

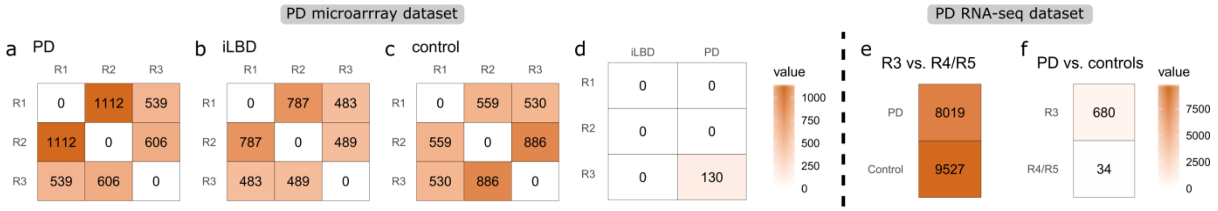


**Supplementary Figure 3 Number of differentially expressed genes in PD microarray and RNA-sequencing datasets.** Differential expression (t-test) was analyzed (a,b,c,e) between regions within groups of individuals and (d, f) between PD or iBLD patients vs. non-demented age-matched controls within regions. Results showed that the expression variation is greater between regions than between groups of individuals.

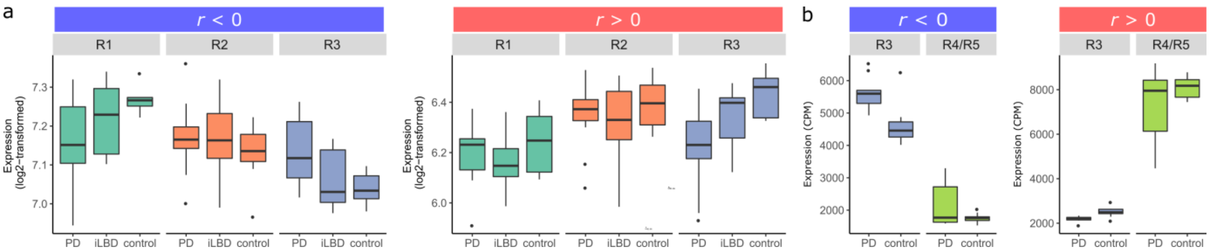


**Supplementary Figure 4 Expression of BRGs in groups of individuals per region in the PD data sets.** Same data is shown in Figure 2f and Figure 2g where samples are plotted per region within groups of individuals (number of samples in Supplementary Table 5 and 7).

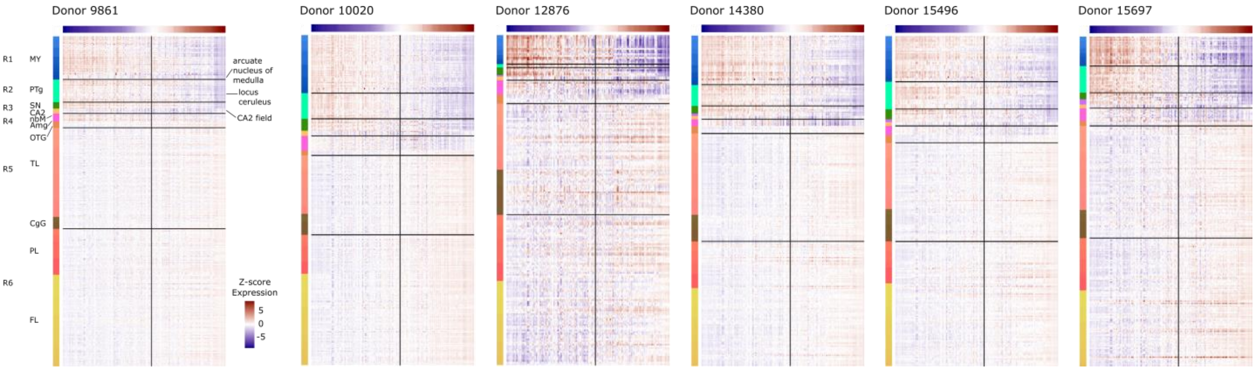


**Supplementary Figure 5 Expression heatmap module eigengenes across Braak region 1-6 for all six donors.** Modules were obtained by hierarchical clustering of genes co-expressed in Braak stage-related regions R1-R6 (rows) with anatomical structures being colored. Eigengenes were sorted based on their correlation with Braak labels (columns). Expression of highly correlated eigengenes either increases from high (red) to low (blue) expression or decreases across regions R1-R6. All modules showed more extreme values (high and low) in R1-R3 compared to R4-R6.

PD microarrray dataset

a R1 vs. R3

| | PD | iLBD | control |
|---|---|---|---|
| neurons | 337 | 133 | 212 |
| astrocytes | 107 | 230 | 225 |
| oligodendrocytes | 273 | 242 | 56 |
| microglia | 242 | 228 | 98 |
| endothelial cells | 261 | 68 | 130 |

b PD vs. controls

| | R3 | R2 | R1 |
|---|---|---|---|
| neurons | 10 | 9 | 0 |
| astrocytes | 95 | 0 | 0 |
| oligodendrocytes | 0 | 0 | 0 |
| microglia | 20 | 0 | 0 |
| endothelial cells | 30 | 0 | 0 |

value: 300, 200, 100, 0

PD RNA-seq dataset

c R3 vs. R4/R5

| | PD | control |
|---|---|---|
| endothelial cells | 194 | 727 |
| microglia | 4 | 569 |
| oligodendrocytes | 471 | 663 |
| astrocytes | 707 | 818 |
| neurons | 33 | 467 |

d PD vs. controls

| | R4/R5 | R3 |
|---|---|---|
| endothelial cells | 0 | 167 |
| microglia | 1 | 11 |
| oligodendrocytes | 0 | 0 |
| astrocytes | 0 | 56 |
| neurons | 0 | 17 |

value: 800, 600, 400, 200, 0

**Supplementary Figure 6 Number of differentially expressed BRGs in the PD datasets after correction for cell-type composition.** Differential expression after cell-type correction with PSEA was analyzed between regions within groups of individuals and between PD patients vs. non-demented age-matched controls within regions. Results showed that the expression variation is greater between regions than between groups of individuals, even after correcting expression differences for five main cell-types.

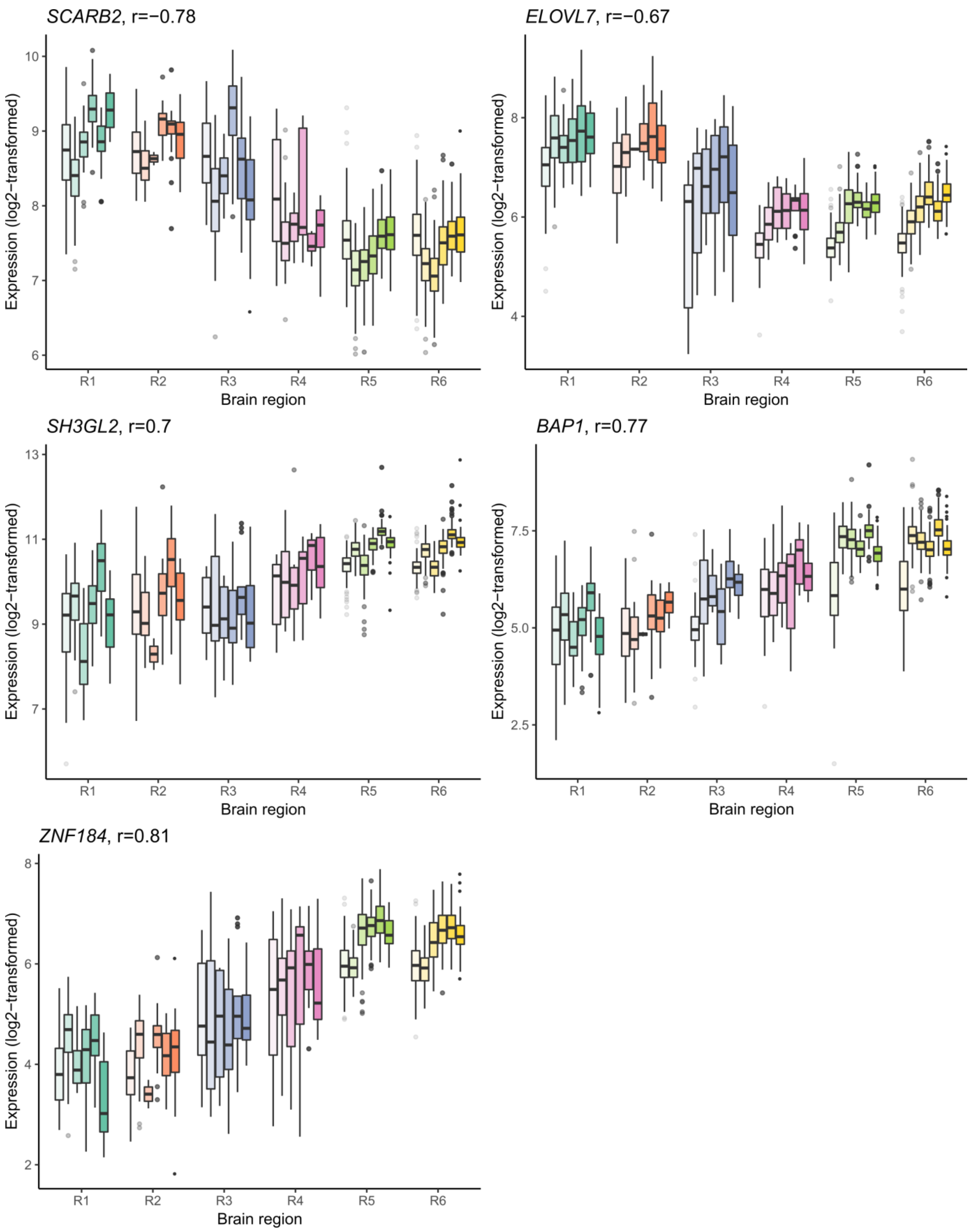


**Supplementary Figure 7 Physiological expression of PD-variant associated genes across regions involved in Braak Lewy body stages.** *SCARB2*, *ELOVL7*, *SH3GL2*, *BAP1*, and *ZNF184* have been associated to PD in gene-wide association studies. Here, we identified expression patterns across regions involved in Braak Lewy body stages of PD (number of samples in Supplementary Table 1).

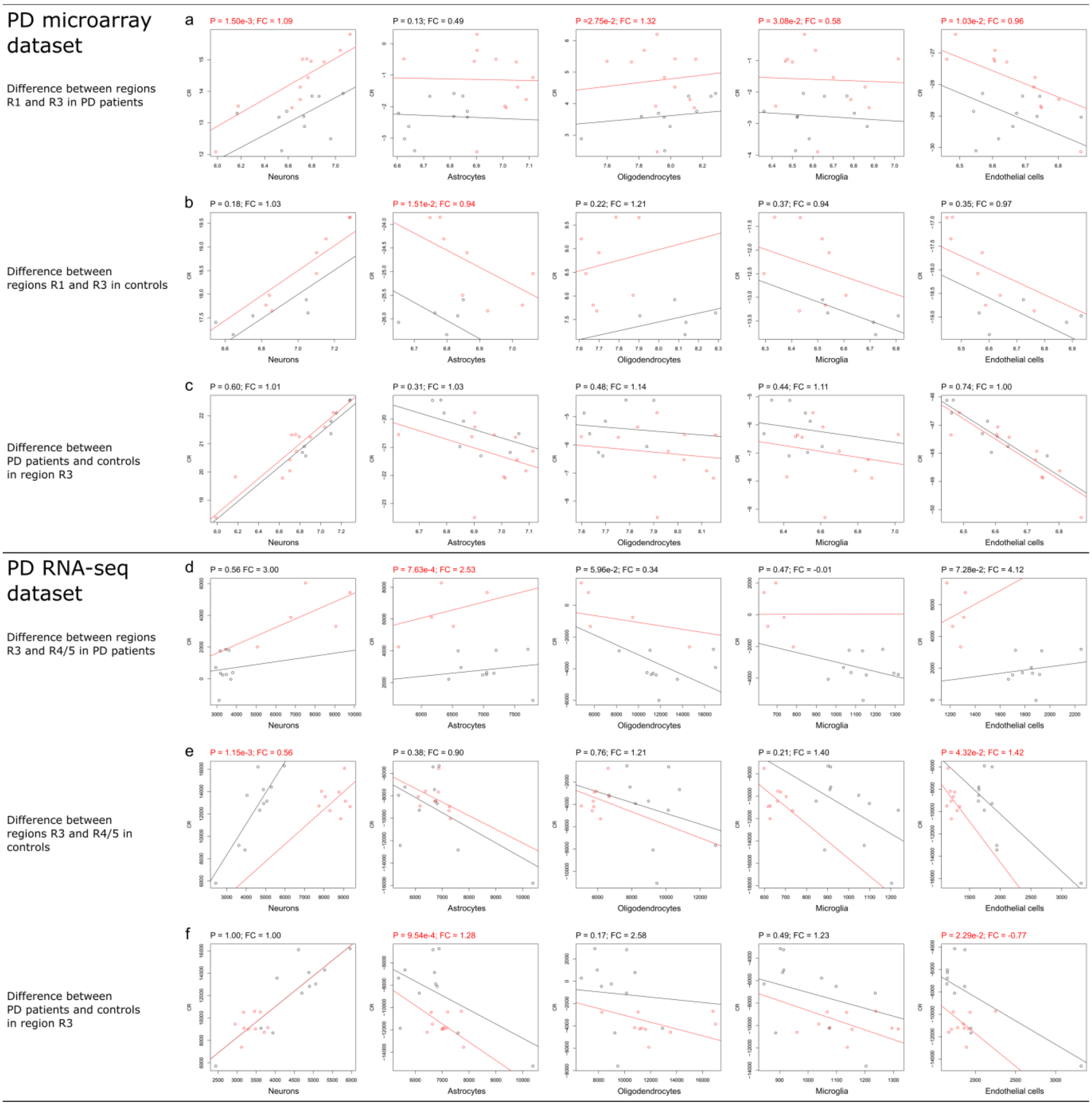


**Supplementary Figure 8 Differential expression of *SNCA* in PD cohorts corrected for cell-type abundance with PSEA.** A microarray and RNA-sequencing dataset of PD cohorts were analyzed for differential expression while correcting for the abundance of neurons, astrocytes, oligodendrocytes, microglia, and endothelial cells based on the average expression of cell-type markers. PSEA analysis was applied between regions as well as between conditions PD and control. Significant BH-corrected P-values are highlighted in red text together with cell-type specific fold-changes (FC; slope change of red line). (a) In microarray data of PD patients, *SNCA* was still significantly differentially expressed between regions R1 (black) and R3 (red) after correcting for neurons, oligodendrocytes, microglia, and endothelial cells. (b) In age-matched controls, *SNCA* was only significantly differentially expressed between R1 and R3 when the change was specific to astrocytes. (c) *SNCA* was not significantly differential expressed between controls (red) and PD patients (black) within region R3 given any of the cell-types. (d) In RNA-seq data of PD patients, *SNCA* was differentially expressed between regions R3 (black) and R4/5 (red) after correcting for astrocytes. (e) In age-matched controls, *SNCA* was differentially expressed between regions R3 (black) and R4/5 (red) after correcting for neurons and endothelial cells. (f) When comparing samples from region R3 between PD patients (red) and controls (black), *SNCA* was only significant given astrocytes and endothelial cells.

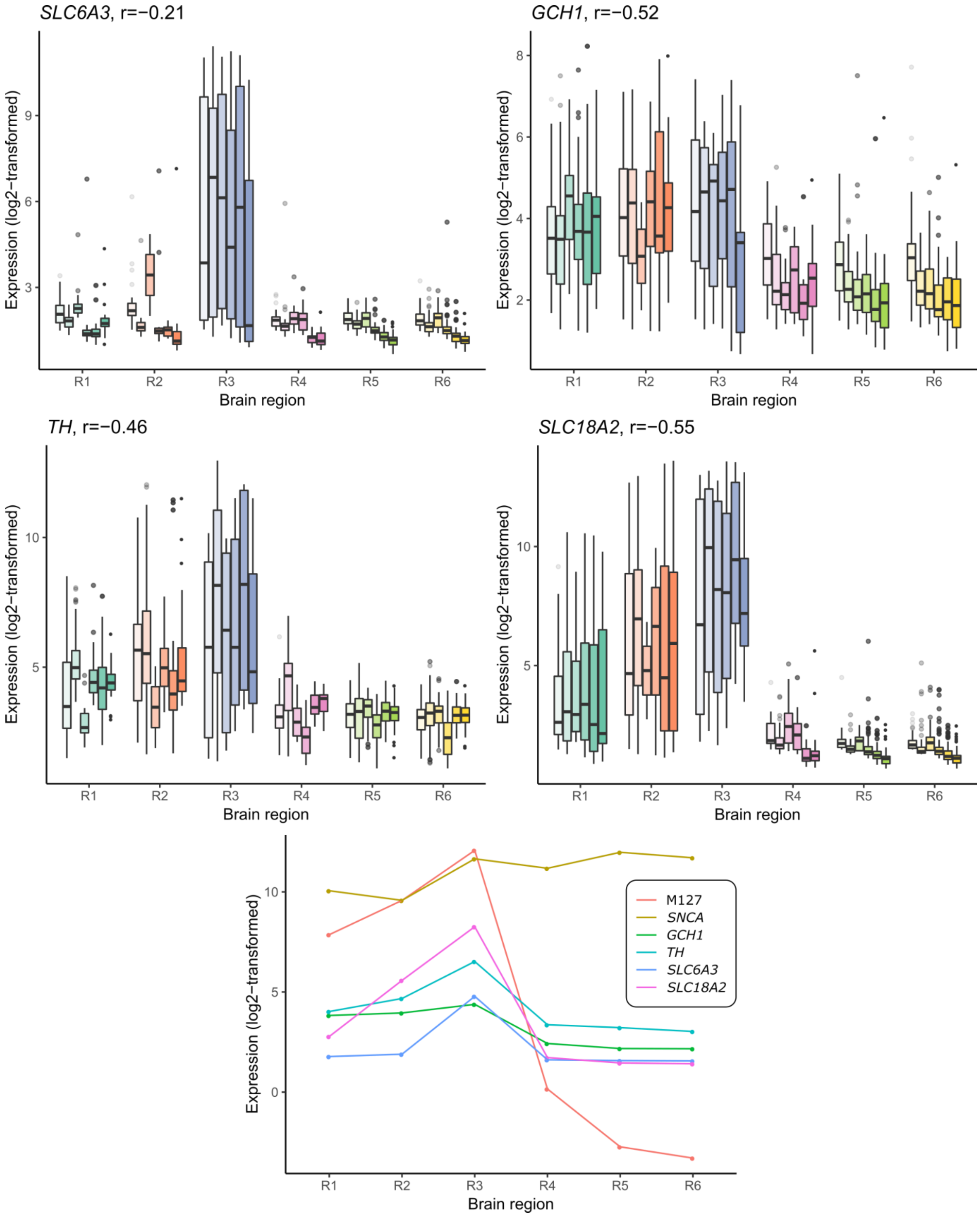


**Supplementary Figure 9 Physiological expression of dopaminergic genes across regions involved in Braak Lewy body stages.** *SLC6A3* (*DAT*), *GCH1*, *TH*, and *SLC18A2* (*VMAT2*) were co-expressed in module M127 which was functionally related to dopamine synthesis. Bottom plot shows the mean of medians across the six donors.

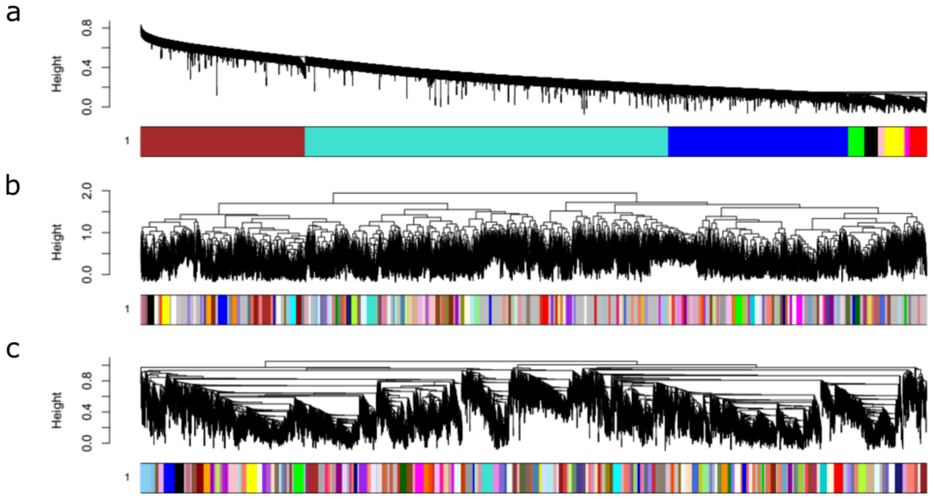


| | Number of modules | Missed genes |
|---|---|---|
| a | 9 | 2 |
| b | 165 | 5745 |
| c | 167 | 1184 |

**Supplementary Figure 10 Hierarchical clustering based on co-expression of genes in Braak regions 1 to 6.** Dendrograms and colored modules obtained with single (A), complete (B) and average linkage (C) are shown together with the number of missed genes and number of modules.
